# Population-specific effect of *Wolbachia* on the cost of fungal infection in spider mites

**DOI:** 10.1101/620310

**Authors:** Flore Zélé, Mustafa Altintaş, Inês Santos, Ibrahim Cakmak, Sara Magalhães

## Abstract

Many studies have revealed the ability of the endosymbiotic bacteria *Wolbachia* to protect its arthropod hosts against diverse pathogens. However, as *Wolbachia* may also increase the susceptibility of its host to infection, predicting the outcome of a particular *Wolbachia*-host-pathogen interaction remains elusive. Yet, understanding such interactions is crucial for disease and pest control strategies. *Tetranychus urticae* spider mites are herbivorous crop pests, causing severe damage on numerous economically important crops. Due to the rapid evolution of pesticide resistance, biological control strategies using generalist entomopathogenic are being developed. However, although spider mites are infected with various *Wolbachia* strains worldwide, whether this endosymbiont protects them from fungi is as yet unknown. Here, we compared the survival of two populations, treated with antibiotics or harbouring different *Wolbachia* strains, after exposure to the fungal biocontrol agents *Metarhizium brunneum* and *Beauveria bassiana*. In one population, *Wolbachia* affected survival in absence of fungi but not in their presence, whereas in the other population *Wolbachia* increased the mortality induced by *B. bassiana*. To control for potential effects of the bacterial community of spider mites, we also compared the susceptibility of two populations naturally uninfected by *Wolbachia*, treated with antibiotics or not. The antibiotic treatment increased the susceptibility of spider mites to *M. brunneum* in one naturally *Wolbachia*-uninfected population, but it had no effect in the other treatments. These results highlight the complexity of within-host pathogens interactions, and the importance of considering the whole bacterial community of arthropods when assessing the effect of *Wolbachia* in a particular system.

## INTRODUCTION

The maternally-inherited bacterium *Wolbachia*, first identified in *Culex pipiens* mosquitoes (Hertig and Wolbach 1924), is to date the best studied and probably the most common endosymbiont of arthropods. Indeed, it is estimated to infect up to 52% of arthropod species (Weinert et al. 2015), a success mainly attributed to its ability to induce various types on reproductive manipulation in hosts to increase the reproductive success of infected females, thereby increasing its own transmission (Werren et al. 2008). Such manipulations include induction of parthenogenetic reproduction of females, feminization of males (i.e. genotypic males develop as phenotypic females), male killing (i.e. infected males die during embryogenesis or late larval instars to the advantage of surviving infected female siblings) and cytoplasmic incompatibility (CI; i.e. modification of male sperm so that females cannot produce offspring unless they are infected with the same strain of *Wolbachia* as their mates). Such ability of *Wolbachia* to rapidly spread within and among host populations (Engelstadter and Hurst 2009), has thus raised growing interests in using it in biocontrol programmes (Bourtzis 2008; Bourtzis et al. 2014; Islam 2007).

Possible *Wolbachia*-based biocontrol strategies include the use of *Wolbachia* as microbial biocontrol agent, for instance to enhance productivity of natural predators and parasites such as parasitoids (e.g. Grenier et al. 1998; Stouthamer 1993; Stouthamer et al. 1999); as a potential gene-drive vehicle for population replacement strategies through cytoplasmic drive (i.e. provides a mechanism for the autonomous spread of desired genes into targeted populations; (Dobson 2003; Dobson et al. 2002; Durvasula et al. 1997; Sinkins and Godfray 2004; Turelli and Hoffmann 1999); or for sterile insect techniques (SIT) to suppress target pest populations by repeated sweeps with infected individuals (Calvitti et al. 2015; Laven 1967; Zhang et al. 2015a; Zhong and Li 2014). Subsequently, the recent discovery of the ability of *Wolbachia* to protects its hosts against a wide array of pathogens, including viruses, protozoan parasites, fungi, or pathogenic bacteria (reviewed in Cook and McGraw 2010) has provided new avenues for the control of vector-borne diseases (reviewed by Iturbe-Ormaetxe et al. 2011; Vavre and Charlat 2012). For instance, deliberate introductions of *Wolbachia* into *Aedes aegypti* mosquito populations are currently being successfully undertaken in several regions worldwide to control dengue virus (e.g. Hoffmann et al. 2014; 2011; Nguyen et al. 2015). However, such ability of *Wolbachia* to interfere with diverse host pathogens may have undesirable effects on biocontrol strategies if, for instance, the bacterium interferes with parasitic biocontrol agents, a possibility that has been largely overlooked. Conversely, natural *Wolbachia* infections in several host species may also increase host susceptibility to parasite infection (e.g. Fytrou et al. 2006; Graham et al. 2012; reviewed in Hughes et al. 2014), raising the possibility that *Wolbachia* could actually facilitate the action of other biocontrol agents. Hence, assessing the effect of *Wolbachia* infection on the efficiency of different strains and/or species of parasitic biocontrol agents should be a prerequisite for the development of efficient and long-lasting control strategies (Zindel et al. 2011).

Spider mite of the genus *Tetranychus* (Acari: Tetranychidae) are ubiquitous major crop pests of c.a. 1100 plant species belonging to more than 140 different plant families (Migeon and Dorkeld 2006-2017). Due to their short generation time and high fecundity, spider mites rapidly develop resistance to pesticides (Van Leeuwen et al. 2010), which led to the development of alternative control strategies such as the use of essential oils or natural enemies (e.g. predators, entomopathogenic bacteria and fungi; (Attia et al. 2013). Among them, entomopathogenic fungi have been successfully used in Integrated Pest Management (IPM) programs, and commercial formulations are currently available to farmers in most parts of the world (Skinner et al. 2014). In particular, fungi such as *Beauveria bassiana, Metarhizium* spp., *Isaria* spp. and *Lecanicillium* spp., have been identified as good candidates for efficient spider mite control (e.g. Bugeme et al. 2008; Chandler et al. 2005; Maniania et al. 2008; Shi et al. 2008; Shin et al. 2017), and their compatibility with other control methods, such as predatory mites (e.g. Dogan et al. 2017; Seiedy 2014; Seiedy et al. 2012; Seiedy et al. 2013; Ullah and Lim 2017; Vergel et al. 2011; Wu et al. 2016) or pesticides (e.g. Irigaray et al. 2003; Klingen and Westrum 2007; Shi et al. 2005) is widely studied. Curiously, however, the interaction between entomopathogenic fungi and bacterial endosymbionts of spider mites has, to our knowledge, never been investigated. This is at odd with the fact that, on the one hand, natural populations of spider mites often carry several maternally-inherited endosymbiotic bacteria with variable prevalence, *Wolbachia* being the most prevalent (Gotoh et al. 2007; Liu et al. 2006; Zélé et al. 2018b; Zhang et al. 2016); and, on the other hand, that *Wolbachia* has been shown to protect *Drosophila melanogaster* hosts against the mortality induced by *B. bassiana* (Panteleev et al. 2007, although no such effect has been found in *D. simulans*; Fytrou et al. 2006).

To examine the effect of the interaction between *Wolbachia* and fungal infection on spider mite survival, we carried out a fully factorial experiment using two naturally *Wolbachia*-infected and two naturally *Wolbachia*-uninfected spider mite populations belonging to the green and the red form of *T. urticae*, and treated or not with antibiotics. We used two generalist entomopathogenic fungi species, *Beauveria bassiana* and *Metarhizium brunneum*, as *Beauveria* and *Metarhizium* spp. are among the most used fungi in commercial production (Vega et al. 2009), with wide geographical and host ranges (Greif and Currah 2007; Gurlek et al. 2018; Meyling and Eilenberg 2007; Rehner 2005; Roberts and Leger 2004). The specific aims of this work were to determine: (i) whether infection with a natural *Wolbachia* strain protects spider mites against fungus-induced mortality, (ii) whether this effect depends on the *Wolbachia* strain and/or other bacteria present in spider mites, and (iii) whether it depends on the fungus species. We then discuss possible mechanisms leading to our results, the importance of considering the whole bacterial community of arthropods when assessing the effect of *Wolbachia* in a particular system, as well as the potential consequences of the presence of *Wolbachia* for the success of spider mite control strategies using entomopathogenic fungi.

## MATERIALS AND METHODS

### Spider mite populations and rearing

Four populations of *Tetranychus urticae* were used in this study, two (AlRo and AMP) belonging to the red form (also referred to as *T. cinnabarinus* by some authors; e.g. Li et al. 2009; Shi et al. 2005; Shi and Feng 2004), and two (DEF and TOM) belonging to the green form. These populations have been collected in the Iberian Peninsula from 2010 to 2017, on different plant species. All the information concerning these populations is summarized in Table S1. After collection, these populations were reared in the laboratory under standard conditions (24±2ºC, 60% RH, 16/8h L/D) at high numbers (c.a. 500-1000 females per population) in insect-proof cages containing bean plants (*Phaseolus vulgaris*, cv. Contender seedlings obtained from Germisem, Oliveira do Hospital, Portugal).

### Endosymbiont infection

Upon collection from the field, the populations AMP (red form) and TOM (green form) were naturally infected by two different strains of *Wolbachia*, although these strains are very closely related, having 1 SNP difference on the sequences of both the fbpA and coxA genes in the multilocus sequence typing system developed by Baldo et al. (2006) for *Wolbachia*. The population AMP is infected by the *Wolbachia* strain ST481 (isolate ‘Turt_B_wUrtAmp’; id: 1858 in the PubMLST *Wolbachia* MLST database available at http://www.pubmlst.org/wolbachia/); and the population TOM is infected by the *Wolbachia* strain ST280 (isolate ‘Turt_B_wUrtTom’; id: 1857). The two other populations, AlRo (red form) and DEF (green form), were naturally uninfected by *Wolbachia* and none of the populations used in this study carried other maternally-inherited bacterial endosymbionts (i.e. *Cardinium, Rickettsia, Spiroplasma, Arsenophonus*) at the time of the experiment, as confirmed by PCR using the methods described in Zélé et al. (2018b; 2018c).

### Antibiotic treatments

Roughly 2 months (i.e. 4 spider mite generations) before the onset of the experiment, a rifampicin solution (0.05%, w/v) was used to treat mites (n=70 adult females initially) from each population for one generation (Gotoh et al. 2005). This allowed to obtain *Wolbachia*-uninfected AMP and TOM populations as well as controls for the antibiotic treatment for the naturally uninfected populations AlRo and DEF. During the treatment, mites were maintained in petri dishes containing bean leaf fragments placed on cotton with the antibiotic solution. After one generation, 100 adult mated daughters from each treated population were transferred in insect-proof cages containing bean plants, in the same laboratory conditions than the untreated populations, and these new populations were allowed to grow for 3 successive generations to avoid potential side effects of antibiotic treatment (Ballard and Melvin 2007; O’Shea and Singh 2015; Zeh et al. 2012). One generation before the onset of the experiment pools of 100 females were taken from each treated population and checked by PCR to confirm that they were uninfected by *Wolbachia* as described in Zélé et al. (2018c).

### Entomopathogenic fungi strains and preparation of inoculum

We used the strains V275 (= Met52, F52, BIPESCO 5) of *Metarhizium brunneum* and UPH-1103 of *Beauveria bassiana*, obtained from Swansea University (UK) and from Siedlce University (Poland), respectively, as they were previously shown to have the potential to suppress *T. urticae* populations (Dogan et al. 2017). The procedures used for fungal growth, inoculum preparation and spider mite infection are similar to that described in Dogan et al. (2017). Briefly, the two fungi were grown on Sabouraud Dextrose Agar (SDA) medium at 25 °C for 2 weeks. Conidia were harvested from sporulating cultures with the aid of a spatula, washed with sterile distilled water and filtered through 4 layers of gauze to remove any hyphae.

### Spider mite infection and survival

The experiment was conducted in a growth chamber under standard conditions (25 ± 2°C, 80% RH, 16/8 h L/D). Roughly 2 weeks prior to the experiment, ca. 100 females were collected from each mass culture and allowed to lay eggs during 4 days on detached bean leaves placed on water-soaked cotton. One day prior to the onset of the experiment, 20 young adult mated females (hence with similar age) were randomly collected from these cohorts and placed on a 9cm^2^ bean leaf disc placed on wet cotton (to ensure the leaf remained hydrated) with the abaxial (underside) surface facing upwards. On the first day of the experiment, the surface of the leaf discs was sprayed using a hand sprayer with 2.5 ml of a spore suspension of *M. brunneum* or *B. bassiana* in 0.03% (v/v) aqueous Tween 20 at 1 × 10^7^ conidia/ml, or, as control, with 0.3% aqueous Tween 20 only. Twelve replicates per treatment and per population were performed within 2 experimental blocks of one day difference (6 replicates of each treatment per block).

### Statistical analysis

Analyses were carried out using the R statistical package (version 3.5.3). The general procedure for building the statistical models was as follows. Survival data were analysed using Cox proportional hazards mixed-effect models (coxme, kinship package). Spider-mite populations (AlRo, AMP, DEF and TOM), antibiotic treatment (treated with rifampicin or not), and infection treatment (sprayed with BB: *Beauveria bassiana*, with MB: *Metarhizium brunneum*, or with Tween 20 only as control) were fitted in as fixed explanatory variables, whereas discs nested within population and block were fitted as random explanatory variables. To explore significant three-way interaction between the three fixed variables, each population was then analyzed separately with the same model structure, except that the variable population was removed from the model. Hazard ratios (HR) were then obtained from these models as an estimate of the difference between the rates of dying (i.e. the instantaneous rate of change in the log number of survivors per unit time; (Crawley 2007) between the untreated controls and the BB or MB treatments for each population.

Because the timing of infection is an important parameter for the fitness of parasites, an additional early measurement of survival, the proportion of dead mites at 3 days post infection (dpi), was obtained from Kaplan–Maier estimates of the survival distribution for each disc. We chose this timing as it is close to (or already above) the median survival upon infection in most of the populations tested, and hence corresponds to a threshold time-point to unravel important differences between treatments. The numbers of dead and alive mites at 3 dpi were computed using the function cbind and analysed with a mixed model glmmadmb procedure (glmmADMB package) with a negative binomial error distribution (family “nbinom1” with a Øµ variance; chosen based on the Akaike information criterium) to correct for overdispersed errors. As above, populations, infection and antibiotic treatment were fitted in as fixed explanatory variables, whereas block was fitted as random explanatory variable.

Maximal models, including all higher-order interactions, were simplified by sequentially eliminating non-significant terms and interactions to establish a minimal model (Crawley, 2007). The significance of the explanatory variables was established using using chi-squared tests (Bolker 2008). The significant chi-squared values given in the text are for the minimal model, whereas non-significant values correspond to those obtained before deletion of the variable from the model.

To further explore significant interactions between infection and antibiotic treatment effects on both HR and mortality at 3 dpi, the two factors were concatenated to fit a single fixed factor containing all treatments levels in the models. Multiple comparisons were then performed using General Linear Hypotheses (glht, package multicomp) with Bonferroni corrections.

## RESULTS

Overall, we found a significant three-way interaction between the effect of the infection by fungi (control females sprayed with Tween 20 only, BB: females sprayed with *B. bassiana*, or MB: females sprayed with *M. brunneum*), the rifampicin treatment, and the population tested, both on the overall survival of spider mites (*X*^*2*^_*6*_=36.16, p<0.0001), and on their mortality at 3 days post-infection (dpi; *X*^*2*^_*6*_=18.33, p=0.005). Indeed, the separate analyses of each population revealed that the survival of females from different populations, naturally infected or uninfected by *Wolbachia*, was not evenly affected by fungal infection and by rifampicin treatment.

### Effect of fungal infection and of antibiotic treatment in the naturally Wolbachia-uninfected population AlRo

In the population AlRo, we found a significant interaction between infection and rifampicin treatment (*X*^*2*^_*2*_=9.53, p=0.009; Fig. 1a), which was due to a stronger mortality induced by *M. brunneum* in rifampicin-treated mites than in untreated mites (z=2.80, p=0.05), while *B. bassiana* induced the same mortality in both rifampicin-treated and untreated mites (z=−1.20, p=1.00 Fig. 1b; Table S2). In both cases, however, *M. brunneum* induced a stronger mortality than *B. bassiana* (MB *vs.* BB: z= 6.72, p<0.0001 and z= 2.81, p=0.05 in rifampicin-treated and untreated mites, respectively). At 3 dpi, however, no significant interaction between infection and rifampicin treatment (*X*^*2*^_*2*_=1.78, p=0.41; Fig. 1c) and no effect of the antibiotics (*X*^*2*^_*1*_=0.90, p=0.34) were found. Only the effect of infection by the two fungi species was found to severely increase the mortality of both rifampicin-treated and untreated mites at this early age of infection (*X*^*2*^_*2*_=135.68, p<0.00014; see also Table S3 for multiple comparisons).

**Figure 1.**
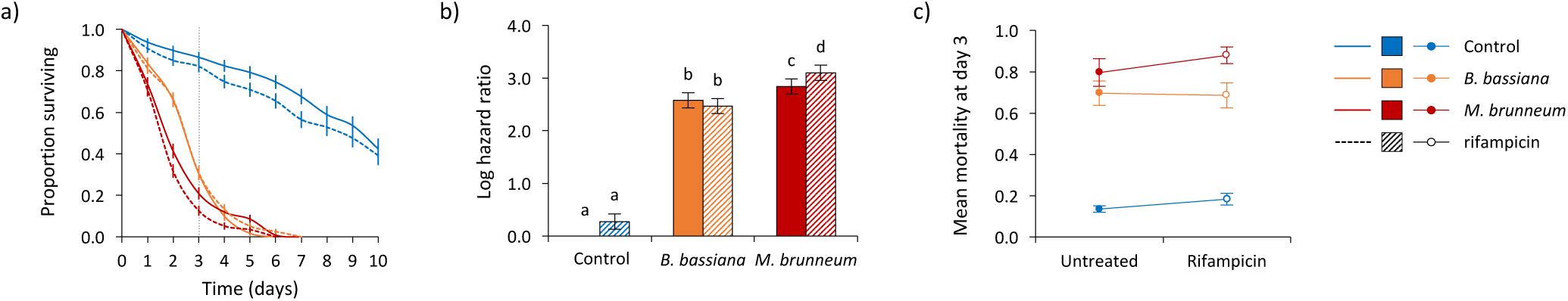
Survival curves (a), relative mortality (b), and survival at 3 dpi (c) of spider mites from the naturally *Wolbachia*-uninfected population AlRo. Adult females were treated (dashed lines, dashed bars and empty circles) or not (solid lines, filled bars and circles) with rifampicin, and sprayed with *B. bassiana* (orange), *M. brunneum* (red), or Tween 20 only as control (blue).

### Effect of fungal infection and of antibiotic treatment in the naturally Wolbachia-uninfected population DEF

In the population DEF, we did not find a significant interaction between infection and rifampicin treatment (*X*^*2*^_*2*_=0.65, p=0.72; Fig. 2a), neither a significant effect of rifampicin treatment (*X*^*2*^_*1*_=0.003, p=0.96), but only a significant effect of fungi infection (*X*^*2*^_*2*_=879.17, p<0.0001). Indeed, both fungi induced the same mortality in both rifampicin-treated and -untreated mites, with an overall stronger effect of *M. brunneum* than of *B. bassiana* (Fig. 2b; see Table S4 for the results of all comparisons). Similarly, at 3 dpi, no significant interaction between infection and rifampicin treatment (*X*^*2*^_*2*_=0.40, p=0.82; Fig. 2c), neither a significant effect of rifampicin treatment (*X*^*2*^_*1*_=0.14, p=0.71) was found. As for the population AlRo, only fungi infection affected the spider-mite survival (*X*^*2*^_*2*_=64.89, p<0.0001; see Table S5 for the results of all comparisons).

**Figure 2.**
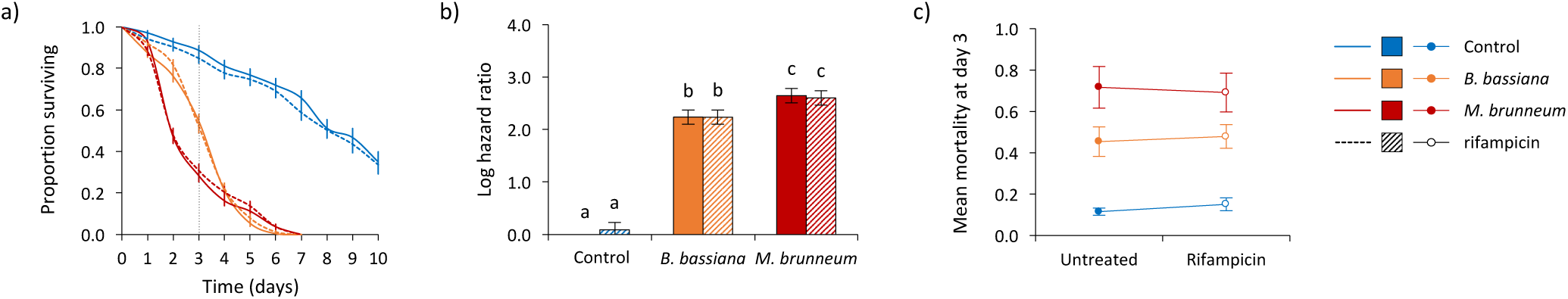
Survival curves (a), relative mortality (b), and survival at 3 dpi (c) of spider mites from the naturally *Wolbachia*-uninfected population DEF. Adult females were treated (dashed lines, dashed bars and empty circles) or not (solid lines, filled bars and circles) with rifampicin, and sprayed with *B. bassiana* (orange), *M. brunneum* (red), or Tween 20 only as control (blue).

### Effect of fungal infection and of antibiotic treatment in the naturally Wolbachia-infected population AMP

In the population AMP, we found a significant interaction between infection and rifampicin treatment (*X*^*2*^_*2*_=26.61, p<0.0001; Fig. 3a), which was due to a lower survival of *Wolbachia*-infected controls compared to uninfected controls (z=−4.92, p<0.0001), while *Wolbachia*-infected and uninfected mites had the same survival upon infection with both fungi species (for *B. bassiana*: z=−0.26, p=1.00; for *M. brunneum*: z=1.88, p=0.55; Fig. 3b and Table S6). Relative to their respective control, both fungi induced higher mortality in rifampicin-treated mites (HR=17.87 and HR=33.39, for BB and MB, respectively) than in *Wolbachia*-infected mites (HR=8.60 and HR=13.21, respectively). A significant interaction between infection and rifampicin treatment was also found at 3 dpi (*X*^*2*^_*2*_=6.5, p=0.04; Fig. 3c). However, this interaction was relatively weak at this time-point, and could not be explained by multiple comparisons between factor levels. Indeed, no differences were found between *Wolbachia*-infected and rifampicin-treated mites when sprayed with Tween 20 only, with *B. bassiana*, or with *M. brunneum* (see Table S7 for the results of all comparisons).

**Figure 3.**
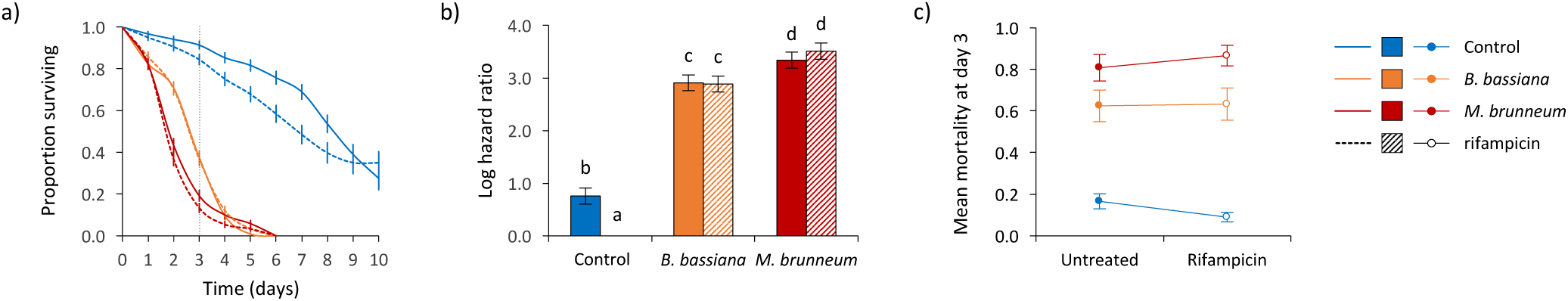
Survival curves (a), relative mortality (b), and survival at 3 dpi (c) of spider mites from the naturally *Wolbachia*-infected population AMP. Adult females were treated (dashed lines, dashed bars and empty circles) or not (solid lines, filled bars and circles) with rifampicin, and sprayed with *B. bassiana* (orange), *M. brunneum* (red), or Tween 20 only as control (blue).

### Effect of fungal infection and of antibiotic treatment in the naturally Wolbachia-infected population TOM

In the population TOM, we also found a significant interaction between infection and rifampicin treatment (*X*^*2*^_*2*_=26.00, p<0.0001; Fig. 4a). In this population, the effect of *B. bassiana* was weaker in rifampicin-treated (HR=3.49) than *Wolbachia*-infected mites (HR=5.53; z=−5.54, p<0.0001; Fig. 4b and Table S8), while *M. brunneum* had the same effect in both rifampincin-treated and untreated mites (HR=6.75 and HR=5.56, respectively; z=1.58, p=1.00). Moreover, whereas both fungi had the same effect on non-treated mites (MB *vs.* BB: z=0.05, p=1.00), *B. bassiana* decreased less the survival of rifampicin-treated treated mites than *M. brunneum* (MB *vs.* BB: z= −6.88, p<0.0001). This effect was even stronger at 3 dpi (interaction between infection and rifampicin: *X*^*2*^_*2*_=15.44, p<0.001). At this time-point, *B. bassiana* induced the same mortality than *M. brunneum* in *Wolbachia*-infected mites (BB *vs.* Control: z=6.24, p<0.0001), but did not affect significantly the survival of rifampicin-treated mites (BB *vs.* Control: z=0.75, p=1.00; Fig. 4c).

**Figure 4.**
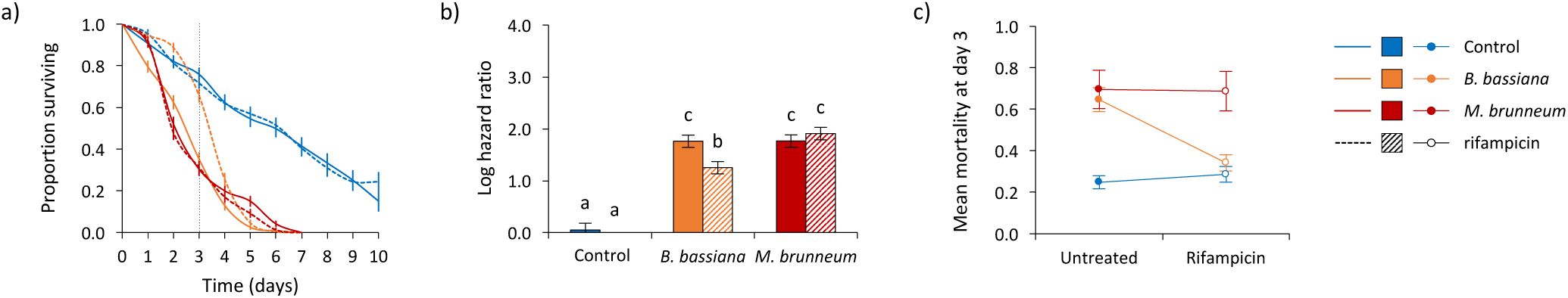
Survival curves (a), relative mortality (b), and survival at 3 dpi (c) of spider mites from the naturally *Wolbachia*-infected population TOM. Adult females were treated (dashed lines, dashed bars and empty circles) or not (solid lines, filled bars and circles) with rifampicin, and sprayed with *B. bassiana* (orange), *M. brunneum* (red), or Tween 20 only as control (blue).

## DISCUSSION

In this study, we found variable effects of infection by *B. bassiana* and *M. brunneum* following antibiotic treatment, depending on the spider mite populations and on whether they were naturally infected by *Wolbachia* or not. Indeed, the mortality induced by both fungi did not differ between *Wolbachia*-infected and uninfected mites in the population AMP, despite *Wolbachia* infection being costly in absence of fungal infection. Similarly, the mortality induced by *M. brunneum* was not affected by *Wolbachia* infection in the population TOM, but that induced by *B. bassiana* increased in presence of *Wolbachia*. These results suggest that Wolbachia may either buffer, or conversely increase, the effect of fungal infection depending on the fungi species, the *Wolbachia* strain and/or the host genetic background. Moreover, in absence of natural *Wolbachia* infection, we found a relatively small effect of the antibiotic treatment on the susceptibility of the mites to infection. Indeed, the antibiotic treatment had no effect on the outcome of infection by fungi, with the exception of a higher mortality in antibiotic-treated mites from the population AlRo when infected with *M. brunneum*. Still, this effect, although significant, is of relatively low amplitude and in the opposite direction than that observed in the *Wolbachia*-infected population TOM following *B. bassiana* infection. This suggests that the effect of *Wolbachia* in the population TOM may not be explained by an alteration of the whole bacterial community in mites following antibiotic treatment. However, because the effect of fungal infection and antibiotic treatment vary between populations independently of the presence of *Wolbachia*, we draw caution on the generalization of such results.

In different arthropod host species, *Wolbachia* may either protect (e.g. Braquart-Varnier et al. 2015; Cattel et al. 2016; Hughes et al. 2011; Kambris et al. 2009; Moreira et al. 2009; Panteleev et al. 2007; Teixeira et al. 2008; Ye et al. 2013), have no effect (e.g. Tortosa et al. 2008; Wong et al. 2011; Zouache et al. 2012), or even increase the susceptibility (e.g. Fytrou et al. 2006; Graham et al. 2012; reviewed in Hughes et al. 2014) of its arthropod hosts to infection depending on the pathogens tested, the *Wolbachia* strain (Chrostek et al. 2013; Martinez et al. 2017; Osborne et al. 2009), but also on the host genetic background, although to a lesser extend (Martinez et al. 2017). In several of these studies the effect of *Wolbachia* on host susceptibility to infection by other pathogens has been assessed following artificial *Wolbachia* infection (e.g. Frentiu et al. 2014; Joubert et al. 2016; Moreira et al. 2009; Walker et al. 2011; Yeap et al. 2011), which prevents a direct alteration of the host bacterial community but may not accurately reflect the effect of natural *Wolbachia* infections. Indeed, novel *Wolbachia* host associations are often costly for hosts (e.g. McGraw et al. 2002), mainly due to the activation of the host immune system following *Wolbachia* infection, which in turn prevents subsequent infections by other pathogens (reviewed in Zug and Hammerstein 2015). Conversely, the effect of natural *Wolbachia* infections on host susceptibility to infection by other pathogens is usually assessed by using antibiotic treatments. However, antibiotics do not affect *Wolbachia* only, but also the entire bacterial community in hosts (e.g. Lehman et al. 2009; Zhu et al. 2018; Zouache et al. 2009), which raises the necessity to assess the potential effect of the antibiotic treatment *per se*.

In *T. truncatus* spider mites, Zhu et al. (2018) showed that an antibiotic treatment (tetracycline hydrochloride during three generations) strongly affects the composition of the bacterial community even after more than 20 generations without antibiotics. In particular, these authors showed that bacteria from different families strongly increased in proportion in tetracycline-treated mites in absence of the Anaplasmataceae (which includes *Wolbachia*). Hence, the lower mortality observed for antibiotic-treated mites following infection by *B. bassiana* in the naturally *Wolbachia*-infected population TOM cannot be unambiguously attributable to *Wolbachia* only. This result could be explained, for instance, by *Wolbachia* outcompeting bacteria that participate to the host homeostasis and immunity (reviewed in Selosse et al. 2014; Shapira 2016; Vavre and Kremer 2014; Weiss and Aksoy 2011), thereby increasing the success of *B. bassiana* infection (i.e., indirect facilitation; Zélé et al. 2018a). In contrast, in the *Wolbachia*-uninfected population AlRo, antibiotic-treated mites have a higher mortality than untreated mites when infected with *M. brunneum*. One possible explanation is that, in absence of natural *Wolbachia* infection, the antibiotic treatment affected differently the bacterial community, potentially eliminating bacteria that interfere with *M. brunneum*.

The apparent facilitation of *B. bassiana* by *Wolbachia* in the TOM population may also be due to *Wolbachia* interacting directly with the host immune system. Indeed, *Wolbachia* has been shown to downregulate autophagy-associated genes in naturally infected hosts, possibly as an immune evasion strategy (Chevalier et al. 2012; Kremer et al. 2009). Under such scenario, the elimination of *Wolbachia* with antibiotics may result in overall higher autophagic processes in the host, to which *B. bassiana* could be susceptible. Moreover, in diverse native hosts, including *T. urticae, Wolbachia* also plays a role in redox homeostasis (Zhang et al. 2015b; reviewed by Zug and Hammerstein 2015). The elimination of *Wolbachia* with antibiotics in coevolved *T. urticae* hosts may thus potentially lead to a disruption of redox homeostasis and higher production of reactive oxygen species (ROS), which are involved in host immunity (e.g. encapsulation, melanisation; (reviewed in Zug and Hammerstein 2015), thereby increasing host resistance to infection. However, all these different scenarios would only explain our results if such mechanisms affect differently the two fungus species and are specific to the *Wolbachia* strain and/or the host population.

As stated above, the host genetic background also plays a major role in determining host susceptibility to infection. First, as recently shown in several spider mite species (Zélé et al. 2019), not all populations (independently of their status of infection by *Wolbachia*) are equally affected by the infection by the two fungi (e.g. the mortality induced by both fungi is stronger in the population DEF than in the population TOM). Second, host susceptibility to infection may also result from G × G interactions with their endosymbionts, as shown, for instance, for *Wolbachia*-mediated protection against viruses across *Drosophila* species (Martinez et al. 2017). Here, the different effects of *Wolbachia* observed in the populations TOM and AMP cannot be unambiguously attributed to the *Wolbachia* strain only, but likely also result from their interaction with the host genetic background. Hence, although further investigations on the respective role of the *Wolbachia* strain and of the host genome, as well as on the composition of the bacterial communities in each of the population tested would be necessary to shed light on the mechanisms involved, these results show that the outcome of infection strongly depends on complex interactions between multiple microorganisms and their host.

In conclusion, our results show variable effects of *Wolbachia* on spider mite susceptibility to fungi-induced mortality using two generalist fungi, *B. bassiana* and *M. brunneum*. To our knowledge, this is the first study investigating the interaction between *Wolbachia* and entomopathogenic fungi on the survival of different spider mite populations within a single full factorial experiment. As *Wolbachia* was found to have either no effect or to increase spider mite susceptibility to fungal infection, these results suggest that it may improve the success of biological control using entomopathogenic fungi. However, these results also highlight the complexity of within-host pathogens interaction, and draw caution on the generalization of the effects of *Wolbachia* as they may vary depending on both the *Wolbachia* strain and the host genetic background. Finally, our findings also point to the importance of considering the whole bacterial community of arthropods when assessing the effect of *Wolbachia* in a particular system.

## Supporting information

Electronic supplementary materials

## AUTHORS’ CONTRIBUTIONS

Experimental conception and design: FZ, SM; maintenance of spider mite populations and plants: IS; acquisition of data: MA; statistical analyses: FZ; paper writing: FZ, SM, with input from all authors. Funding: IC, SM. All authors have read and approved the final version of the manuscript.

## ACKNOWLEDGMENTS

We thank Catarina Pinto and João Alpedrinha for their help in some parts of the experiment, as well as Marta Palma for technical support. We also thank all members of the SM lab for useful discussions and suggestions. This work was funded by an FCT-Tubitak agreement (FCT-TUBITAK/0001/2014 and TUBITAK TOVAG 115O610) to IC and SM, and by Adnan Menderes University Research Foundation (ZRF-17055) to IC. FZ was funded through an FCT Post-Doc fellowship (SFRH/BPD/125020/2016). Funding agencies did not participate in the design or analysis of experiments. We declare that we do not have any conflict of interest.

