## Supplementary material for "Population-specific effect of *Wolbachia* on the cost of fungal infection in spider mites": Electronic supplementary materials

**Table S1.** Populations of spider mites used in the experiment. Mites were collected in Portugal (P) and Spain (S), and were naturally infected, or not, by *Wolbachia*. The absence of other maternally-inherited endosymbionts (*Cardinium*, *Rickettsia, Spiroplasma, Arsenophonus*) in these populations was confirmed by PCR before the onset of the experiment (using methods described in [1, 2]).

| Name | Date | Host plant | Location | Coordinates | *Wolbachia* infection | Ref |
|---|---|---|---|---|---|---|
| AlRo | 09/11/2013 | *Rosa spp.* | Almería (S) | 36.855725, -2.320374 | no | [2] |
| DEF | 26/04/2017 | *Solanum lycopersicum* | Alvalade, Lisbon (P) | 38.75515, -9.14685 | no | - |
| AMP | 18/11/2013 | *Datura stramonium* | Aldeia da Mata Pequena (P) | 38.534363, -9.191163 | yes (ST481)*[1] | [2] |
| TOM | --/05/2010 | *Solanum lycopersicum* | Carregado (P) | 39.078962,-8.993656 | yes (ST280)*[2] | [3] |

***[1] Isolate 'Turt_B_wUrtTom' - id: 1857, *Wolbachia* strain ST280.** This strain has been first identified as wTurt_2 from three different populations of *T. urticae* in China [4].

***[2] Isolate 'Turt_B_wUrtAmp' - id: 1858, *Wolbachia* strain ST481.** This is a new strain, very similar to the strain ST219 (they differ by 1 SNP on the fbpA gene: allele 444 instead of allele 4), which belongs to the supergroup B of *Wolbachia* and was found in China [5].

**Table S2.** Results of multiple comparisons (with Bonferroni correction) between hazard ratios obtained for the naturally *Wolbachia*-uninfected population AlRo sprayed or not with fungi (BB: *Beauveria bassiana*; MB: *Metarhizium brunneum*; Control: Tween 20 only) and treated or not with antibiotics (rif: rifampicin-treated; nt: untreated).

| Factor 1 | Factor 2 | Estimate | Std. Error | z value | Pr(>\|z\|) |
|---|---|---|---|---|---|
| Control_rif | Control_nt | 0.272 | 0.146 | 1.869 | 0.554 |
| BB_rif | BB_nt | -0.110 | 0.092 | -1.200 | 1.000 |
| MB_ rif | MB_nt | 0.259 | 0.093 | 2.802 | 0.046* |
| BB_ nt | Control_nt | 2.578 | 0.144 | 17.931 | < 2e-16*** |
| MB_nt | Control_nt | 2.840 | 0.144 | 19.718 | < 2e-16*** |
| MB_ nt | BB_nt | 0.262 | 0.093 | 2.810 | 0.045* |
| BB_ rif | Control_rif | 2.195 | 0.134 | 16.345 | < 2e-16*** |
| MB_rif | Control_rif | 2.827 | 0.139 | 20.410 | < 2e-16*** |
| MB_ rif | BB_rif | 0.632 | 0.094 | 6.718 | 1.66E-10*** |

**Table S3.** Results of multiple comparisons (with Bonferroni correction) between mortality at 3 dpi of the naturally *Wolbachia*-uninfected population AlRo sprayed or not with fungi infection (BB: *Beauveria bassiana*; MB: *Metarhizium brunneum*; Control: Tween 80 only) and treated or not with antibiotics (rif: rifampicin-treated; nt: untreated).

| Factor 1 | Factor 2 | Estimate | Std. Error | z value | Pr(>\|z\|) |
|---|---|---|---|---|---|
| Control_rif | Control_nt | 0.327 | 0.236 | 1.387 | 1.000 |
| BB_rif | BB_nt | -0.012 | 0.247 | -0.049 | 1.000 |
| MB_ rif | MB_nt | 0.085 | 0.175 | 0.489 | 1.000 |
| BB_ nt | Control_nt | 1.684 | 0.196 | 8.607 | < 2e-16*** |
| MB_nt | Control_nt | 1.818 | 0.194 | 9.386 | < 2e-16*** |
| MB_ nt | BB_nt | 0.134 | 0.232 | 0.579 | 1.000 |
| BB_ rif | Control_rif | 1.345 | 0.291 | 4.614 | 3.55E-05*** |
| MB_rif | Control_rif | 1.576 | 0.276 | 5.702 | 1.06E-07*** |
| MB_ rif | BB_rif | 0.232 | 0.202 | 1.145 | 1.000 |

**Table S4.** Results of multiple comparisons (with Bonferroni correction) between hazard ratios obtained for the naturally *Wolbachia*-uninfected population DEF sprayed or not with fungi (BB: *Beauveria bassiana*; MB: *Metarhizium brunneum*; Control: Tween 20 only) and treated or not with antibiotics (rif: rifampicin-treated; nt: untreated).

| Factor 1 | Factor 2 | Estimate | Std. Error | z value | Pr(>\|z\|) |
|---|---|---|---|---|---|
| Control_rif | Control_nt | 0.091 | 0.141 | 0.648 | 1.000 |
| BB_rif | BB_nt | -0.001 | 0.092 | -0.015 | 1.000 |
| MB_ rif | MB_nt | -0.044 | 0.091 | -0.487 | 1.000 |
| BB_ nt | Control_nt | 2.238 | 0.136 | 16.435 | < 2e-16*** |
| MB_nt | Control_nt | 2.647 | 0.138 | 19.240 | < 2e-16*** |
| MB_ nt | BB_nt | 0.410 | 0.096 | 4.246 | 1.96E-04*** |
| BB_ rif | Control_rif | 2.145 | 0.132 | 16.220 | < 2e-16*** |
| MB_rif | Control_rif | 2.512 | 0.134 | 18.685 | < 2e-16*** |
| MB_ rif | BB_rif | 0.367 | 0.095 | 3.845 | 0.001** |

**Table S5.** Results of multiple comparisons (with Bonferroni correction) between mortality at 3 dpi of the naturally *Wolbachia*-uninfected population DEF sprayed or not with fungi infection (BB: *Beauveria bassiana*; MB: *Metarhizium brunneum*; Control: Tween 80 only) and treated or not with antibiotics (rif: rifampicin-treated; nt: untreated).

| Factor 1 | Factor 2 | Estimate | Std. Error | z value | Pr(>\|z\|) |
|---|---|---|---|---|---|
| Control_rif | Control_nt | 0.177 | 0.328 | 0.539 | 1.000 |
| BB_rif | BB_nt | 0.089 | 0.368 | 0.241 | 1.000 |
| MB_ rif | MB_nt | -0.020 | 0.258 | -0.077 | 1.000 |
| BB_ nt | Control_nt | 1.255 | 0.277 | 4.525 | 5.44E-05*** |
| MB_nt | Control_nt | 1.738 | 0.265 | 6.550 | 5.17E-10*** |
| MB_ nt | BB_nt | 0.482 | 0.339 | 1.423 | 1.000 |
| BB_ rif | Control_rif | 1.167 | 0.416 | 2.806 | 0.045* |
| MB_rif | Control_rif | 1.541 | 0.393 | 3.925 | 0.001*** |
| MB_ rif | BB_rif | 0.374 | 0.304 | 1.231 | 1.000 |

**Table S6.** Results of multiple comparisons (with Bonferroni correction) between hazard ratios obtained for the naturally *Wolbachia*-infected population AMP sprayed or not with fungi (BB: *Beauveria bassiana*; MB: *Metarhizium brunneum*; Control: Tween 20 only) and treated or not with antibiotics (rif: rifampicin-treated; nt: untreated).

| Factor 1 | Factor 2 | Estimate | Std. Error | z value | Pr(>\|z\|) |
|---|---|---|---|---|---|
| Control_rif | Control_nt | -0.756 | 0.154 | -4.916 | 7.93E-06*** |
| BB_rif | BB_nt | -0.024 | 0.091 | -0.258 | 1.000 |
| MB_ rif | MB_nt | 0.172 | 0.092 | 1.875 | 0.547 |
| BB_ nt | Control_nt | 2.151 | 0.136 | 15.819 | < 2e-16*** |
| MB_nt | Control_nt | 2.581 | 0.136 | 18.921 | < 2e-16*** |
| MB_ nt | BB_nt | 0.430 | 0.093 | 4.599 | 3.81E-05*** |
| BB_ rif | Control_rif | 2.883 | 0.150 | 19.195 | < 2e-16*** |
| MB_rif | Control_rif | 3.508 | 0.154 | 22.828 | < 2e-16*** |
| MB_ rif | BB_rif | 0.625 | 0.093 | 6.702 | 1.85E-10*** |

**Table S7.** Results of multiple comparisons (with Bonferroni correction) between mortality at 3 dpi of the naturally *Wolbachia*-infected population AMP sprayed or not with fungi infection (BB: *Beauveria bassiana*; MB: *Metarhizium brunneum*; Control: Tween 80 only) and treated or not with antibiotics (rif: rifampicin-treated; nt: untreated).

| Factor 1 | Factor 2 | Estimate | Std. Error | z value | Pr(>\|z\|) |
|---|---|---|---|---|---|
| Control_rif | Control_nt | -0.644 | 0.270 | -2.390 | 0.152 |
| BB_rif | BB_nt | 0.013 | 0.235 | 0.056 | 1.000 |
| MB_ rif | MB_nt | 0.070 | 0.170 | 0.411 | 1.000 |
| BB_ nt | Control_nt | 1.322 | 0.178 | 7.424 | 1.03E-12*** |
| MB_nt | Control_nt | 1.579 | 0.174 | 9.088 | < 2e-16*** |
| MB_ nt | BB_nt | 0.257 | 0.219 | 1.175 | 1.000 |
| BB_ rif | Control_rif | 1.979 | 0.316 | 6.258 | 3.50E-09*** |
| MB_rif | Control_rif | 2.293 | 0.305 | 7.529 | 4.58E-13*** |
| MB_ rif | BB_rif | 0.314 | 0.195 | 1.608 | 0.970 |

**Table S8.** Results of multiple comparisons (with Bonferroni correction) between hazard ratios obtained for the naturally *Wolbachia*-infected population TOM sprayed or not with fungi (BB: *Beauveria bassiana*; MB: *Metarhizium brunneum*; Control: Tween 20 only) and treated or not with antibiotics (rif: rifampicin-treated; nt: untreated).

| Factor 1 | Factor 2 | Estimate | Std. Error | z value | Pr(>\|z\|) |
|---|---|---|---|---|---|
| Control_rif | Control_nt | -0.050 | 0.130 | -0.385 | 1.000 |
| BB_rif | BB_nt | -0.510 | 0.092 | -5.539 | 2.74E-07*** |
| MB_ rif | MB_nt | 0.145 | 0.092 | 1.579 | 1.000 |
| BB_ nt | Control_nt | 1.711 | 0.120 | 14.237 | < 2e-16*** |
| MB_nt | Control_nt | 1.715 | 0.120 | 14.302 | < 2e-16*** |
| MB_ nt | BB_nt | 0.004 | 0.093 | 0.045 | 1.000 |
| BB_ rif | Control_rif | 1.250 | 0.118 | 10.603 | < 2e-16*** |
| MB_rif | Control_rif | 1.909 | 0.121 | 15.839 | < 2e-16*** |
| MB_ rif | BB_rif | 0.659 | 0.096 | 6.878 | 5.48E-11*** |

**Table S9.** Results of multiple comparisons (with Bonferroni correction) between mortality at 3 dpi of the naturally *Wolbachia*-infected population TOM sprayed or not with fungi infection (BB: *Beauveria bassiana*; MB: *Metarhizium brunneum*; Control: Tween 80 only) and treated or not with antibiotics (rif: rifampicin-treated; nt: untreated).

| Factor 1 | Factor 2 | Estimate | Std. Error | z value | Pr(>\|z\|) |
|---|---|---|---|---|---|
| Control_rif | Control_nt | 0.125 | 0.177 | 0.706 | 1.000 |
| BB_rif | BB_nt | -0.637 | 0.234 | -2.717 | 0.059. |
| MB_ rif | MB_nt | -0.012 | 0.175 | -0.069 | 1.000 |
| BB_ nt | Control_nt | 0.949 | 0.152 | 6.239 | 3.96E-09*** |
| MB_nt | Control_nt | 1.024 | 0.151 | 6.798 | 9.56E-11*** |
| MB_ nt | BB_nt | 0.075 | 0.200 | 0.374 | 1.000 |
| BB_ rif | Control_rif | 0.187 | 0.250 | 0.751 | 1.000 |
| MB_rif | Control_rif | 0.887 | 0.220 | 4.033 | 4.95E-04*** |
| MB_ rif | BB_rif | 0.699 | 0.205 | 3.414 | 0.006** |
